# Rapid, noninvasive, and unsupervised detection of sleep/wake using piezoelectric monitoring for pharmacological studies in narcoleptic mice

**DOI:** 10.1101/226522

**Authors:** Sarah Wurts Black, Jessica D. Sun, Alex Laihsu, Nikki Kimura, Pamela Santiago, Kevin D. Donohue, Bruce F. O’Hara, Ross Bersot, Paul S. Humphries

## Abstract

**Background:** Assessment of sleep/wake by electroencephalography (EEG) and electromyography (EMG) is invasive, resource intensive, and not amenable to rapid screening at scale for drug discovery. In the preclinical development of therapeutics for narcolepsy, efficacy tests are hindered by the lack of a non-EEG/EMG based translational test of symptom severity. The current methods study offers proof-of-principle that PiezoSleep (noninvasive, unsupervised piezoelectric monitoring of gross body movement, together with respiration patterns during behavioral quiescence), can be used to determine sleep/wake as applicable to the development of wake-promoting therapeutics. First, the translational wake-maintenance score (WMS, the ratio of time during the first half of the dark period spent in long wake bouts to short sleep bouts) of the PiezoSleep narcolepsy screen was introduced as a means by which to rank narcoleptic orexin/ataxin-3 mice and wild type mice by sleep/wake fragmentation severity. Accuracy of the WMS to detect narcoleptic phenotypes were determined in genotype-confirmed orexin/ataxin-3 mice and wild type colony mates. The WMS was used to identify the most highly symptomatic mice for resource-intensive EEG/EMG studies for further analysis of specific arousal states. Second, PiezoSleep was demonstrated for use in high-throughput screening of wake-promoting compounds using modafinil in orexin/ataxin-3 and wild type mice.

**Results:** The WMS detected a narcoleptic phenotype with 89% sensitivity, 92% specificity and 98% positive predictive value. A 15-fold difference in WMS differentiated wild type littermates from the most severely affected orexin/ataxin-3 mice. Follow-up EEG/EMG study indicated 82% of the orexin/ataxin-3 mice with the lowest wake-maintenance scores met or exceeded the cataplexy-occurrence threshold (≥ 3 bouts) for inclusion in therapeutic efficacy studies. In the PiezoSleep dose-response study, the ED_50_ for wake-promotion by modafinil was approximately 50 mg/kg in both genotypes. Using unsupervised piezoelectric monitoring, the efficacy of wake-promoting compounds can be determined in a 5-arm study with 60 mice in less than one week—a fraction of the time compared to EEG/EMG studies.

**Conclusions:** The WMS on the PiezoSleep narcolepsy screen quantifies the inability to sustain wakefulness and provides an accurate measure of the narcoleptic phenotype in mice. PiezoSleep offers rapid, scalable assessment of sleep/wake for high-throughput screening in drug discovery.

## Background

Pharmacological studies with arousal state endpoints are typically invasive, resource intensive, and time consuming. Rodent polysomnography requires surgical implantation of electrodes, recovery time, adaptation to sleep implant hardware, and manual scoring of arousal states for the highly accurate detection of sleep microstructure. Using standard electroencephalography (EEG) and electromyography (EMG), wakefulness is defined as low amplitude, desynchronous activity in the EEG across a broad frequency range with variable EMG activity. Sleep in rodents is classified as either rapid-eye-movement (REM) sleep, with pronounced activity in the EEG theta band (4.5-9.5 Hz) and atonia in the EMG, or non-REM (NREM) sleep, with slow, synchronous activity in the EEG delta band (0.5 to 4.5 Hz) and low, steady EMG tone. Some of the burden of rodent polysomnography has been relieved by the development of automated methods of scoring EEG/EMG records. However, the expediency and objectivity value of these methods is diminished for those that require a subset of manually scored samples from individual animals to train the algorithm (1-4). Unsupervised, automated scoring algorithms have addressed these limitations of machine learning with varying degrees of accuracy (72-96% agreement) compared to standard manual scoring (5-8). Nevertheless, all EEG/EMG based systems of arousal state monitoring are hindered by the time required for surgery, recovery, and in many cases, the repeated-measures experimental designs used for studies in which a small number of animals are used. There is a need in drug discovery for a fully automated method of sleep/wake staging that can scale to a sample size that is amenable to high-powered, between-subject designs for the rapid screening of compounds.

The use of non-invasive proxy-measures of sleep/wake, such as actigraphy, plethysmography, or heartbeat rate correlates of arousal states instead of traditional polysomnography, can expedite sleep/wake determination in rodents (9-14). In particular, the use of a highly sensitive motion detector (piezoelectric sensor) on the cage floor to monitor gross motor activity, together with breathing patterns during periods of behavioral quiescence, yields unsupervised sleep/wake detection with 90% accuracy vs. manual EEG scoring (15). This piezoelectric sleep monitoring system (PiezoSleep) has been used to characterize daily sleep/wake amounts and patterns in a transgenic mouse model of Alzheimer’s disease (16) and in 180 knockout lines as part of a large-scale phenotyping program (17).

Transgenic *orexin/ataxin-3* (Atax) mice model the human sleep disorder narcolepsy with high fidelity. Like their human counterparts, Atax mice experience degeneration of wake-promoting orexin (also known as hypocretin) neurons which leads to cataplexy (a sudden, reversible loss of muscle tone and REM-sleep like EEG) and/or increased sleep propensity at times of day when wakefulness would otherwise predominate (18). Although Atax mice exhibit cataplexy that is readily identifiable according to EEG/EMG-dependent consensus criteria (19), high inter-individual variability in cataplexy occurrence presents a challenge to sort mice for inclusion in studies with cataplexy reduction as an endpoint (20). Currently, there is no prescreening method prior to EEG/EMG studies for sorting mice that are highly symptomatic for narcolepsy. In a previous study, Atax mice were monitored with piezoelectric detection of movement and heartrate for a few hours during the light period, but the algorithm for autoscoring was only 73% accurate against manual EEG scoring (14) and missed the time of day in which cataplexy and sleep/wake fragmentation occurs most frequently (20).

While cataplexy is pathognomonic of narcolepsy, excessive daytime sleepiness (EDS) is the most problematic symptom in patients with narcolepsy (20). The EDS leads to an inability of patients with narcolepsy to sustain long periods of wakefulness during the Maintenance of Wakefulness Test (MWT) and to fall asleep with short latency on the Multiple Sleep Latency Test (MSLT). Both tests are indicators of symptom severity; the MSLT is used in the diagnosis of narcolepsy and the MWT is used to assess treatment efficacy (21). Modafinil (MOD) is a first-line choice of treatment for EDS in patients and promotes EEG/EMG-defined wakefulness in narcoleptic mice but is ineffective in cataplexy reduction (21, 22). Although neural mechanisms (23) and response to treatment (24, 25) may differ between cataplexy and sleep/wake fragmentation, a link between these symptoms is suggested by the finding that patients with severe cataplexy are the least able to stay awake on the MWT (26). Drug discovery efforts aimed at finding improved therapeutics for narcolepsy would benefit from a translational assay in mice similar to the MSLT and MWT that is rapid, scalable, and not dependent upon polysomnography nor sleep deprivation protocols (27).

In the current report, we demonstrated the proof of principle that the PiezoSleep system can be applied to address two problems in the preclinical development of wake-promoting therapeutics for narcolepsy. First, we addressed the need for a non-EEG/EMG based translational test of sleepiness severity that can be used for subject stratification by introducing the wake-maintenance score (WMS)—a measure of sleep/wake fragmentation determined by the PiezoSleep narcolepsy screen. The subset of Atax mice that were identified with the lowest WMS were subjected to follow-up polysomnography to quantify cataplexy. Second, we conducted a dose-response study with MOD to demonstrate that PiezoSleep can fill the need for a non-invasive, rapid means to quantify sleep time and consolidation at a scale amenable to screening wake-promoting therapeutics in drug discovery.

## Methods

All experimental procedures were approved by the Institutional Animal Care and Use Committee at Reset Therapeutics and were conducted in accordance with the principles set forth in the *Guide for Care and Use of Laboratory Animals of the National Institute of Health*.

### Animals

Male and female hemizygous transgenic B6.Cg-Tg(HCRT-MJD)1Stak/J mice (Atax) and wild type (WT) littermates (JAX stock #023418, Jackson Labs, Bar Harbor, ME) were used (18). Separate cohorts were used for immunohistochemistry, development of the narcolepsy phenotyping assay and PiezoSleep narcolepsy screen, calculation of the receiver operator characteristic (ROC) curve for the WMS, EEG/EMG and video monitoring, and pharmacology. Male C57BL/6J mice (JAX stock #000664, Jackson Labs, Bar Harbor, ME) were also used in pharmacological studies. Male CD-1 mice (Charles River Laboratories, Hollister, CA), 7-8 weeks old, were used in pharmacokinetic studies. Animals were individually housed, except for CD-1 mice which were housed 3 per cage, at 22-25 °C, 30-70% relative humidity, and LD12:12 with *ad libitum* access to food, water and nesting material.

### Immunohistochemistry

All histological procedures and cell counts were performed at Jackson Laboratories (Bar Harbor, ME). Male and female Atax (*N* = 8 and 5, respectively) and WT littermates (*N* = 8 males and 6 females), aged 16.9 ± 0.1 and 28.5 ± 0.8 g were euthanized by pentobarbital overdose and transcardially perfused with 0.4 % paraformaldehyde. Brains were harvested, postfixed overnight, and cryo sectioned at 40 μm from approximately -1.23 to -1.91 relative to Bregma in coronal sections. After blocking in normal donkey serum (5% in 0.3% triton in PBS), free-floating brain sections were incubated overnight at room temperature with goat anti-donkey orexin A antibody (SC-8070, Santa Cruz Biotechnology, CA) at 1:2000 dilution. After washing with phosphate-buffered saline, sections were incubated for 2 hours at room temperature with biotinylated donkey anti-goat (Jackson Immunoresearch, West Grove, PA) at 1:500 dilution in blocking solution. After further washes, sections were developed with the avidin-biotin-peroxidase system using diaminobenzidine as the chromogen (Vector Laboratories, Burlingame, CA). Sections were mounted onto slides, and anatomically matched sections between brains that represented the anterior, mid, and posterior regions of the orexin field were identified for manual cell counting. Counts were performed bilaterally on these sections at 20x magnification under light microscopy. Only somata with complete and intact cell membranes were included in the analysis.

### Piezoelectric monitoring

The narcoleptic phenotype of Atax mice and their sleep/wake responses in pharmacological studies were determined by piezoelectric recording of gross body movement and breath rate (PiezoSleep version 2.11, Signal Solutions, Lexington, KY). The PiezoSleep apparatus consisted of 60 individual polycarbonate cages, each of which contained a polyvinylidene difluoride piezo sensor beneath a thin (0.025 cm) plastic shield. The sensor covered the entire area of the cage floor (232.3 cm^2^) and was connected to a single channel tube amplifier (RoMo version 1.2, Signal Solutions, Lexington, KY). Each amplifier fed into a digital acquisition device connected to a computer for data collection and sleep/wake determination as previously described (9, 15, 28). Briefly, signals were amplified and filtered between 0.5 and 10 Hz and analog-to-digital converted at 120 Hz sampling rate. The frequency, amplitude, and peak spectral energy of the signal were evaluated in 2 second increments over a tapered, 8 second window to automatically calculate a sleep/wake decision statistic by simple linear discriminant analysis. Signals were classified as sleep if they exhibited a periodicity with a consistent, low relative amplitude at a frequency in the typical breathing range (1-4 Hz). Wakefulness was associated with high amplitude signals with variable frequency and broad spectral energy. Signal features related to these properties were extracted and combined to obtain a decision statistic for each interval. Over time, the decision statistics per increment accumulated in a bimodal distribution. The saddle point of the distribution was determined using an adaptive threshold for the classification of sleep and wake. Bouts of contiguous intervals of sleep or wake were defined to terminate when an interval, equal to the minimum bout length of 30 s, included less than 50% of the targeted state.

Piezoelectric experiments began by loading mice into individual PiezoSleep cages that contained standard rodent bedding (diamond dry cellulose approximately 1 cm thick) with food and nesting material from each mouse’s home cage. Mice were permitted at least 24 h of acclimation before experimental data were sampled. For development of the narcolepsy phenotype assay, after the acclimation period, Atax mice (*N* = 31, aged 15.7 ± 0.3 weeks, 29.4 ± 0.4 g) and their WT littermates (*N* = 17, aged 14.8 ± 0.5, 29.5 ± 0.4 g) were recorded undisturbed for 36 h beginning at zeitgeber time (ZT)12 (21:00). Hourly sleep percentages, sleep bout duration (SBD) and wake bout duration (WBD) means per h, and histograms of the percent time spent asleep or awake in bouts from 0.5 to ≥ 64 min long were calculated using SleepStats version 2.11 (Signal Solutions, Lexington, KY). Based on the circadian distribution of SBD and WBD means per h, and on the increased propensity for cataplexy during the early dark period (20), sleep and wake data from ZT12-18 were used to differentiate mice based on their ability to maintain long WBD at that time of day. Scatterplots of time spent in short SBD (< 8 min) vs. long WBD (≥ 32 min) were used to screen Atax mice that were highly symptomatic for murine narcolepsy (i.e., compared with other Atax mice, they spent more than the mean amount of time in short SBD and less than the mean amount of time in long WBD.)

All mice were assigned a WMS based on their ratio of time in long WBD:short SBD during ZT12-18 for the PiezoSleep narcolepsy screen. The optimal threshold for the WMS to detect a narcoleptic phenotype with high sensitivity and specificity was determined by a ROC curve in which the WMS cutoff was adjusted from 0.25-50 on data from all available cohorts (*N* = cohorts) of genotyped Atax (*N* = 148) and wild-type colony mates (*N* = 36).

### EEG/EMG & video monitoring

A separate cohort of male Atax mice (*N* = 37, aged 11.6 weeks, 26.6 ± 0.4 g) was subjected to the PiezoSleep narcolepsy screen to identify the individuals that were the most highly symptomatic for narcolepsy and thus the best candidates for EEG and EMG studies. The 16 Atax mice with the lowest WMS were surgically implanted at 16.9 ± 0.2 weeks of age (30.2 ± 0.7 g) with transmitters (F20-EET, Data Sciences International, St. Paul, MN) for EEG and EMG recording as previously described (29). Mice were anesthetized with isoflurane and transmitters were placed IP along the midline. Biopotential leads were routed subcutaneously to the head, and EMG leads were positioned bilaterally through the nuchal muscles. Cranial holes were drilled 1 mm anterior to bregma and 1 mm lateral to midline, and contralaterally, 2 mm posterior to bregma and 2 mm lateral to midline. EEG leads were placed subchranially over the dura and attached to the skull with cyanoacrylate and dental acrylic. Analgesia was managed with prophylactic ketoprofen (5 mg/kg, SC) followed by buprenorphine (0.1 mg/kg, SC) upon emergence from anesthesia. Ketoprofen (5 mg/kg, SC, QD) was continued for 3 days postsurgery. Mice received postoperative thermal support for 24 h and hydration (0.7 mL lactated Ringer’s solution, SC and dietary recovery gel) for 3 days. Mice were permitted 3 weeks postsurgical recovery, which included 2 weeks adaptation to running wheels and the light-controlled (LD12:12 with ZT0 = 03:00), sound-attenuated and ventilated recording chamber.

Physiological and video-recorded behavioral data were simultaneously acquired with Ponemah version 5.2 (Data Sciences International, St. Paul, MN). Electrophysiological data were sampled at 500 Hz and digital videos were recorded at 15 frames per second (640 x 480 pixels) using Noldus Media Recorder (Leesburg, VA). Recordings from mice in their home cages (with nesting material and running wheels) began at ZT6 and continued for the following 30 h. Data were manually scored using NeuroScore version 3.2.1 (Data Sciences International, St. Paul, MN) by scorers trained with ≥ 96% interrater reliability by an expert scorer. Data were classified in 10-second epochs as wakefulness, wheel-running during wakefulness, NREM sleep, REM sleep, or cataplexy. Criteria for cataplexy were ≥ 10 seconds of EMG atonia, theta-dominated EEG, and video-confirmed behavioral immobility preceded by ≥ 40 seconds of wakefulness (19). Bouts of each arousal state were defined to terminate with the occurrence of a single epoch of another state. Of the 16 implanted mice, 11 were included in the analysis (aged 19.9 ± 0.3 weeks at recording), 2 did not survive surgical recovery, and 3 had corrupted data files or unscorable EEG signals.

### Compound formulation

Modafinil (MOD, 2-[(diphenylmethyl)sulfinyl]acetamide) was purchased from Focus Synthesis (San Diego, CA) for dosing in CD-1 and WT and from Advanced ChemBlocks, Inc. (Burlingame, CA) for dosing in Atax mice because MOD was no longer available from Focus Synthesis. For pharmacokinetic studies, MOD was formulated in 1.25% Hydroxypropyl Methylcellulose / 0.25% Methylcellulose in Water for Injection, which formed a homogenous suspension. For efficacy studies, after MOD was weighed, premade vehicle (20% propylene glycol, 25% polyethylene glycol 400 in water for injection) was added at a volume appropriate for the highest concentration, followed by overnight sonication, which produced a clear solution. Serial dilutions were created and re-sonicated approximately 6 h prior to dosing. All dosing solutions were formulated within 24 h prior to dosing.

### Pharmacokinetic studies

Male CD-1 mice were acclimated for a minimum of 3 days prior to drug administration. Modafinil was administered as 10, 30, 50, or 100 mg/kg via oral gavage (PO) to CD-1 mice at ZT0 to ZT1. Blood and tissue samples were collected from three mice per time point at pre-dose, 15, 30, 60, 90, 180, 360, 720, and 1440 min following oral gavage from the CD-1 mice. Blood (500 μL) was obtained via cardiac puncture into Microtainer Blood Collection Tubes with K_2_EDTA (BD Biosciences, San Jose, CA), and gently mixed. Blood was immediately centrifuged, and the resulting plasma was frozen on dry ice then stored at -80 °C until it was analyzed. Entire brain from each animal was harvested, weighed, frozen on dry ice, and stored at -80 °C until it was analyzed. The concentrations of MOD were determined by a liquid chromatography – mass tandem spectrometry (LC-MS/MS) method (Quintara Discovery, Hayward, CA) following the standard protocol. Briefly, Brain samples were homogenized in two volumes of ice cold water. Then 20 μL of each plasma or brain homogenate sample were processed by protein precipitation with methanol: acetonitrile (5:95 v:v) containing internal standard, verapamil. Extracts were analyzed by LC-MS/MS using positive electrospray ionization under the multiple-reaction-monitoring mode for transitions m/z 274->167 (MOD) and m/z 455-> 165 (verapamil). The lower limit of quantitation was 1.8 nM for plasma and 5.5 nM for tissues. Pharmacokinetic parameters of MOD were generated by non-compartmental analysis with Phoenix™ WinNonlin^®^ software (Pharsight Corporation, St. Louis, MO).

### Efficacy studies

In a between-subjects design, male C57BL/6 mice (WT, 15.7 ± 0.1 weeks, 28.5 ± 0.2 g, *N* = 12/group) and Atax mice (16.4 ± 0.7 weeks, 30.0 ± 0.7 g, *N* = 9-10/group) were dosed separately in two pharmacological studies. A between-subjects design was chosen to ensure mice were drug naïve at the time of dosing. Mice were loaded into PiezoSleep cages for 42 h acclimation, followed by 29 h baseline recording prior to dosing at ZT5 (14:00), the time of day in which wake-promoting/sleep-inhibiting effects would be expected to be most pronounced because baseline sleep is typically high then. Syringes were filled immediately prior to dosing, which was completed within a 15-minute window. After pharmacokinetic studies (Table 1) and based on efficacy in previous reports (22, 30), mice received either MOD (10, 30, 50, 100 mg/kg, 10 mL/kg) or vehicle orally and were recorded for 44 h post dosing. Mice were undisturbed during the recording period except for dosing. Groups were balanced across dosing order, dosing personnel, and cage location per shelf and rack. Atax mice had previously been screened in the narcolepsy phenotype assay, and highly symptomatic mice were balanced across dosing groups.

**Table 1.** Pharmacokinetic data after oral modafinil at ZT0 in CD-1 mice

| Dose<br>(mg/kg) | C<sub>max</sub> (μM) |  | T<sub>1/2</sub> (h) |  | MRT (h) |  | AUC (μM·h) |  |
| --- | --- | --- | --- | --- | --- | --- | --- | --- |
|  | Plasma | Brain | Plasma | Brain | Plasma | Brain | Plasma | Brain |
| 10 | 14.2 ± 0.83 | 4.13 ± 0.89 | 1.1 | 1.4 | 1.8 | 1.5 | 22.5 | 7.3 |
| 30 | 76.1 ± 11 | 15.4 ± 3.1 | 0.9 | 1.2 | 1.7 | 1.5 | 131.3 | 34.5 |
| 50 | 80.1 ± 12 | 33.7 ± 8.4 | 0.8 | 1.2 | 2.1 | 1.9 | 185.5 | 72.4 |
| 100 | 306 ± 50 | 76.5 ± 6.3 | 1.0 | 1.1 | 2.4 | 2.7 | 958.5 | 294.9 |

C<sub>max</sub>: maximum concentration, mean ± standard deviation
T<sub>1/2</sub>: half-life
MRT: mean residence time
AUC: area under the curve

### Data analysis and statistics

Statistics were performed using GraphPad Prism version 7.03 (La Jolla, CA). Unless otherwise noted, all data are presented as the mean ± S.E.M. Orexin cell counts were compared between male and female Atax and WT mice using two-way analysis of variance (ANOVA). In phenotyping studies, sleep time and mean SBD and WBD were compared between Atax and WT mice using two-way mixed-model ANOVA on factors “genotype” (between subjects) and “hour” (within subjects). Histograms representing the distribution of time spent in various SBD and WBD were evaluated using two-way mixed-model ANOVA on factors “genotype” (between subjects) and “bout duration bin” (within subjects). Pearson correlations were used to evaluate the day-to-day variability in individual mice for measures of sleep/wake fragmentation. Wake-maintenance scores were compared between genotypes using two-tailed, unpaired *t* test. For evaluation of the WMS to accurately predict a narcoleptic phenotype using a ROC curve, sensitivity was calculated as the proportion of Atax mice that had a positive WMS for a narcoleptic phenotype out of all Atax genotyped mice. Specificity was calculated as the proportion of WT mice that had a negative WMS for a narcoleptic phenotype out of all mice genotyped as wild type. The positive predictive value of the WMS was determined as the proportion of Atax mice that scored positive for the narcoleptic phenotype out of all positively scored Atax and WT mice. The negative predictive value was calculated as the proportion of WT mice that had a negative WMS out of all WT and Atax mice that scored negative for the narcoleptic phenotype. Comparisons of wake time measurements via piezoelectric monitoring vs. EEG/EMG with video recording over 24 h were made within subjects using two-way repeated-measures (RM) ANOVA, and for 24 h totals using two-tailed, paired *t* test. Pharmacological data were assessed within genotypes by two-way mixed-model ANOVA on between-subjects factor “drug condition” and within-subjects factor “hour” or “bout duration bin.” The maximum sleep deficit was calculated as the minimal value in the cumulative change from baseline in sleep % time during the 12 h post dosing. Consolidation of wakefulness after dosing was measured as the change from baseline in the % time spent in long wake bouts ≥ 8 min. Sleep deficits and consolidated wakefulness were compared between genotypes after dosing using two-way ANOVA. For each analysis, when ANOVA indicated significance (α = 0.05), contrasts between relevant factor levels (vs. WT or vehicle, as appropriate) were detected *post hoc* with Bonferroni’s multiple comparisons test. The total sample size (*N* = 97) for the pharmacological study permitted sufficient power (0.8) for the detection of an effect size of 36% at *P* < 0.05 (computed using WebPower <u>https://webpower.psychstat.org/wiki/</u>).

## Results

### Histology

To confirm orexin cell loss from Atax mice taken out of cryopreservation at JAX, orexin cell counts were compared between the transgenics and their WT littermates at an adult age in which they would be used for sleep studies. Representative images (Figure 1) of the tuberal region of the hypothalamus and cell counts (Table 2) indicated that, as expected, Atax mice had fewer cells that stained positive for orexin A. There were no sex differences in the number of orexin neurons that were sampled, *F*(1, 23) = 3.58, *n.s*. The extent of orexin neurodegeneration was consistent with that observed in a different colony of Atax mice (29).

**Table 2.**
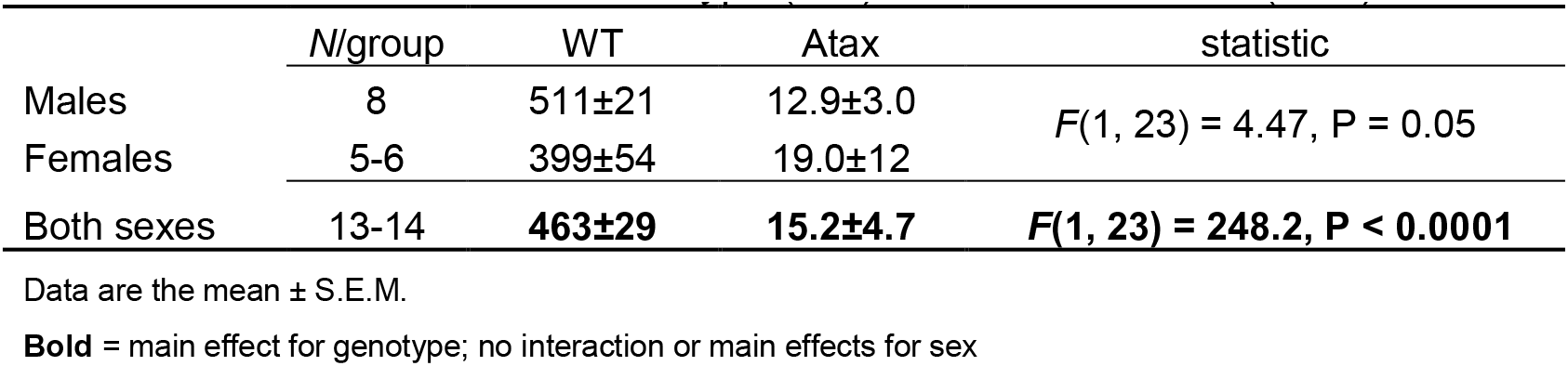
Orexin cell counts in wild type (WT) and *orexin/ataxin-3* (Atax) mice

**Figure 1.**
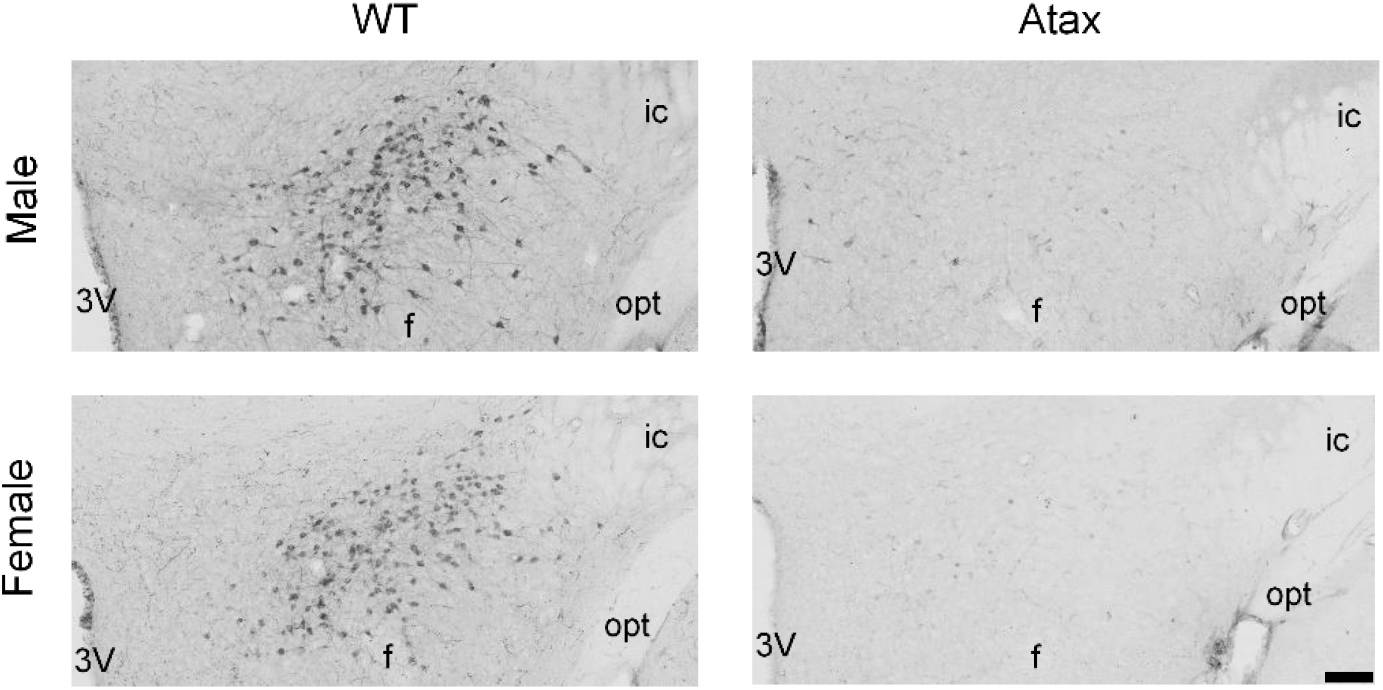
**Loss of orexin neurons in orexin/ataxin-3 (Atax) mice.** Representative images of Orexin A+ cells in the posterior lateral hypothalamus of male **(top)** and female **(bottom)** wild type littermates **(left)** and Atax mice **(right)**. Images taken at 10x magnification (bar = 100 μM), approximately -1.5 mm from Bregma; fornix (f), internal capsule (ic), optic tract (opt), third ventricle (3V).

### Phenotyping of narcolepsy model

Piezoelectric monitoring of sleep/wake was used to evaluate the phenotype of Atax mice and to sort mice by the severity of murine narcolepsy symptoms. Compared to WT littermates, Atax mice slept more at a few time points during the dark period, particularly during the last hour prior to lights-on (Figure 2A), *F*(36, 1656) = 4.81, *P* < 0.0001. However, the total amount of sleep per 24 h was the same between genotypes (WT: 45.5 ± 0.8 % vs. Atax: 45.8 ± 1.0 %), *t*(46) = 0.03, *n.s*., as expected. Genotype differences were observed in sleep and wake bout duration measures. Atax mice exhibited shorter mean SBD per h across the day (Figure 2B, *F*(36, 1649) = 4.65, *P* < 0.0001) and shorter mean WBD during the dark period (Figure 2C, *F*(36, 1656) = 8.62, *P* < 0.0001).

**Figure 2.**
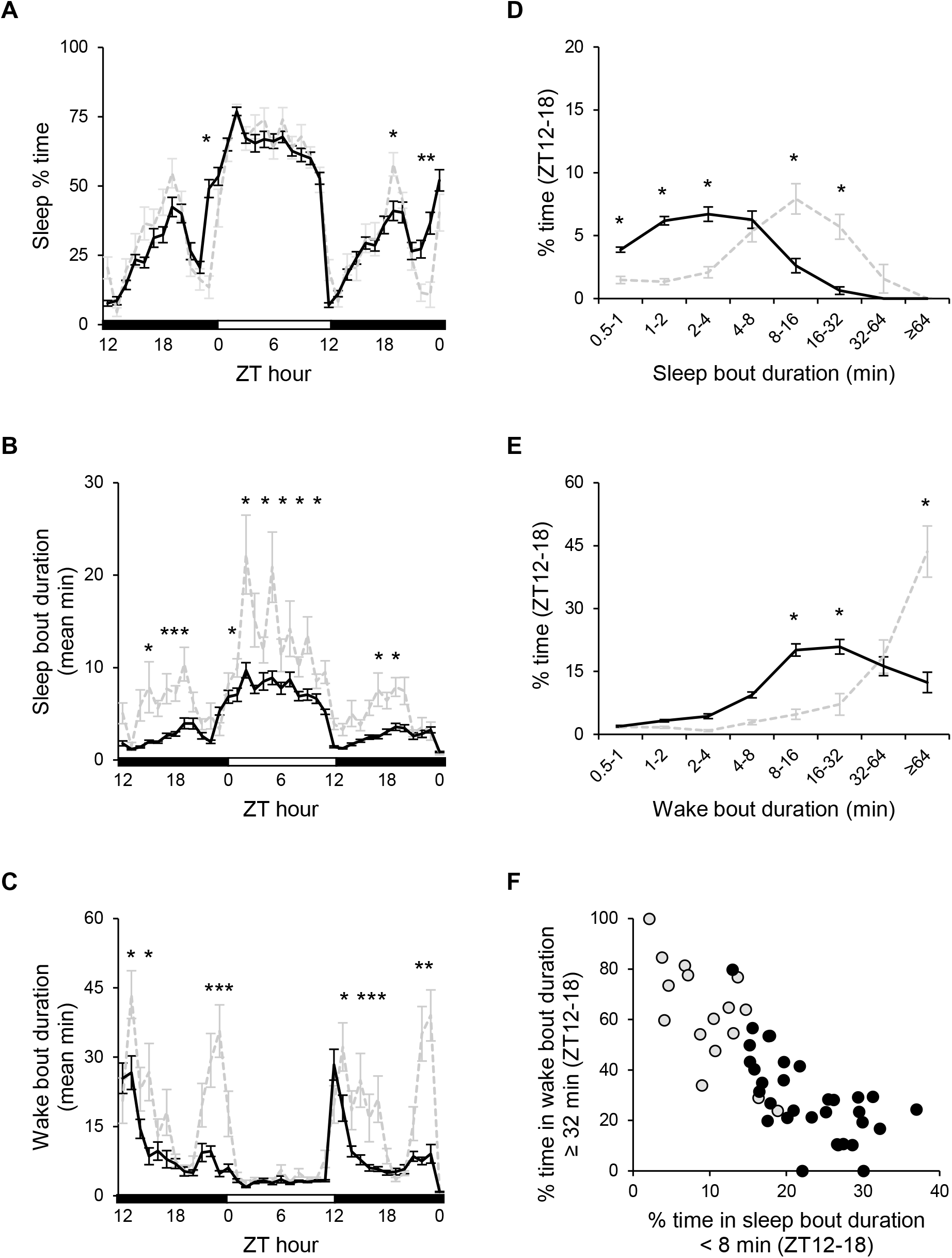
**The PiezoSleep narcolepsy phenotyping assay.** Determination of narcoleptic phenotype in male orexin/ataxin-3 (Atax, *N* = 31) mice **(black)** vs. wild-type (WT, *N* = 17) littermates **(gray, dashed lines)** using PiezoSleep. Sleep % per h **(A)** and mean bout durations per h of sleep **(B)** and wake **(C)** over the 36 h recording period. Percentage of time during the first half of the dark period (zeitgeber time (ZT)12-18) spent asleep **(D)** and awake **(E)** in bout durations lasting from 0.5 to ≥ 64 min. Percentage of time spent in short sleep bouts (< 8 min) vs. long wake bouts (≥ 32 min) during the first half of the dark period per individual mouse **(F)**. Two-way mixed-model analysis of variance with Bonferroni’s multiple comparisons: * P < 0.05 vs. WT.

Because differences in both mean SBD and WBD were particularly robust during the first half of the dark period, and because that time of day is associated with an increased probability of cataplexy occurrence (20), ZT12-18 was chosen as the focus for determination of narcolepsy symptom severity. The percentage of time spent in sleep bouts that were 0.5 to 4 min long was greater in Atax than for WT mice, while WT mice spent more time than Atax mice in sleep bouts that were 8 to 32 min long (Figure 2D), *F*(7, 368) = 24.31, *P* < 0.0001. Compared to WT littermates, Atax mice spent more time in wake bouts that were 8-32 min long and less time in the longest wake bouts of ≥ 64 min duration (Figure 2E), *F*(7, 368) = 25.74, *P* < 0.0001. The percentage of time spent in short SBD (< 8 min) vs. time spent in long WBD (≥ 32 min) were plotted in Figure 2E. Atax mice clustered on the scatterplot in a manner indicative of more time spent in short sleep bouts and less time in long wake bouts than their WT littermates. The ratio of time spent in long WBD to short SBD determined the WMS and was smaller in Atax mice (1.5 ± 0.2) than in WT (9.9 ± 2.7), *t*(46) = 4.19, *P* < 0.0001.

The day-to-day variation in PiezoSleep indices of sleep/wake fragmentation was assessed by correlative comparison of data collected from ZT12-18 on subsequent days. Measures of percent time spent in short SBD, percent time in long WBD, and WMS taken on the two different days highly correlated within individual mice between days (Figure 3).

**Figure 3.**
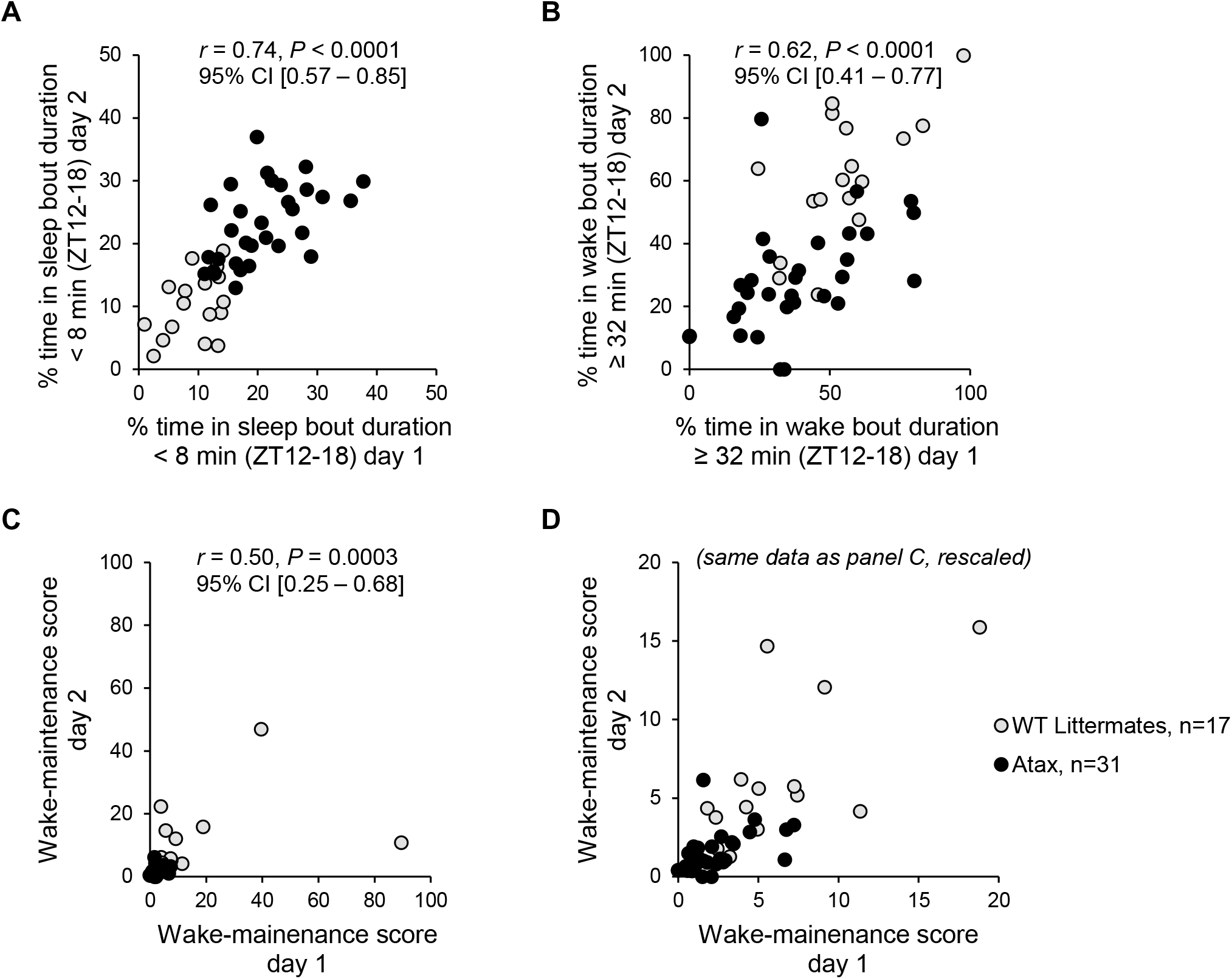
**Reliability of PiezoSleep measures of sleep/wake fragmentation across recording days.** Pearson correlations (with 95% confidence intervals) were used to compare data collected on subsequent days in male orexin/ataxin-3 (Atax, *N* = 31) mice **(black)** and wild-type (WT, *N* = 17) littermates **(gray)**. Data were collected during the first half of the dark period (zeitgeber time (ZT)12-18) for evaluation of the percentage of time spent in short sleep bouts (< 8 min) **(A)**, long wake bouts (≥ 32 min) **(B)**, and wake-maintenance scores (ratio of time in long wake bouts to short sleep bouts) **(C)**. Wake-maintenance scores were rescaled without outliers for clarity **(D)**.

The accuracy of the WMS to detect a narcoleptic phenotype was evaluated using a ROC curve in which the threshold for defining the narcoleptic phenotype was adjusted from 0.25 to 50 on data from mice genotyped as either Atax or WT (Figure 4). Based on the ROC curve, the optimal cutoff for the WMS was 3.25 to detect murine narcolepsy with 89% sensitivity (95% CI [82-93%]), 92% specificity (95% CI [76-98%]), 98% positive predictive value (95% CI [93-99%]), and 66% negative predictive value (95% CI [51-78%]).

**Figure 4.**
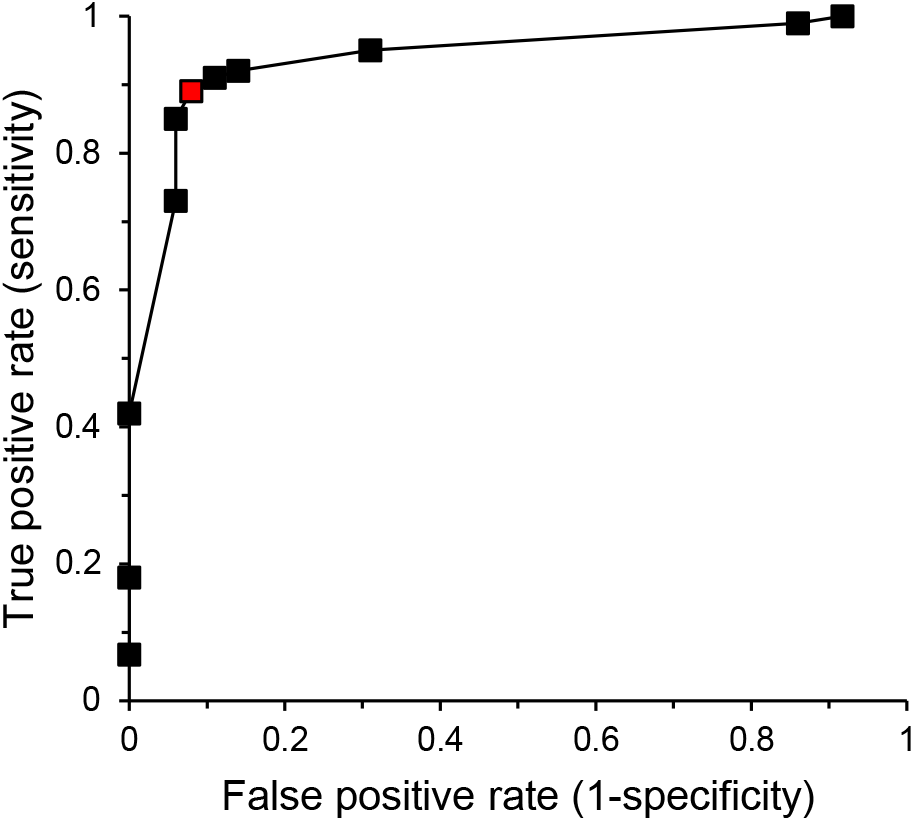
**Receiver operator characteristic (ROC) curve for the wake-maintenance score (WMS) on the PiezoSleep narcolepsy screen.** Accuracy of the WMS (ratio of time spent in wake bouts ≥32 min to sleep bouts < 8 min during the first half of the dark period) to detect a narcoleptic phenotype in genotype-confirmed orexin/ataxin-3 mice (*N* = 148) vs. wild type colony mates (*N* = 36) as the threshold for the WMS was increased from 0.25 to 50. A WMS of 3.25 (red) was determined to be optimal to detect narcolepsy with 89% sensitivity (95% CI [82-93%]), 92% specificity (95% CI [76-98%]), 98% positive predictive value (95% CI [93-99%]), and 66% negative predictive value (95% CI [51-78%]).

The WMS was used to sort individual Atax mice by murine narcolepsy symptom severity to prioritize EEG/EMG instrumentation on the mice most likely to demonstrate cataplexy. Using the PiezoSleep narcolepsy screen on a separate cohort of Atax mice, 16 mice were identified as “highly symptomatic” for murine narcolepsy, with lower WMS (0.68 ± 0.1) than the remaining Atax littermates (3.15 ± 0.4), *t*(35) = 5.17, *P* < 0.0001, and were implanted for EEG/EMG monitoring. The WMS of the 11 mice that were ultimately included in the EEG/EMG analysis (0.67 ± 0.1) were representative of the original 16 animals.

The percentage of time spent awake per hour across 24 h was compared between piezoelectric recordings from the phenotype screen and EEG/EMG with video monitoring that was performed 8 weeks later (Figure 5A). This comparison of non-simultaneous recordings from different environments was not intended to validate the accuracy of the piezoelectric sleep/wake detection against polysomnography, as has previously been done (9, 15), but rather was intended to assess the predictive consistency between the two methods under different use cases. The pattern of the daily distribution of wakefulness appeared to be similar between sleep/wake recording methods, but piezoelectric determination yielded slightly more wakefulness per 24 h (53.3 ± 0.9%) than EEG/EMG with video monitoring (49.7 ± 1.3%), *t*(20) = 2.31, *P* = 0.03). However, the only time point in which the PiezoSleep recordings differed from the EEG/EMG recordings was in the hour immediately prior to lights on, *F*(23, 230) = 29.3, *P* < 0.0001.

**Figure 5.**
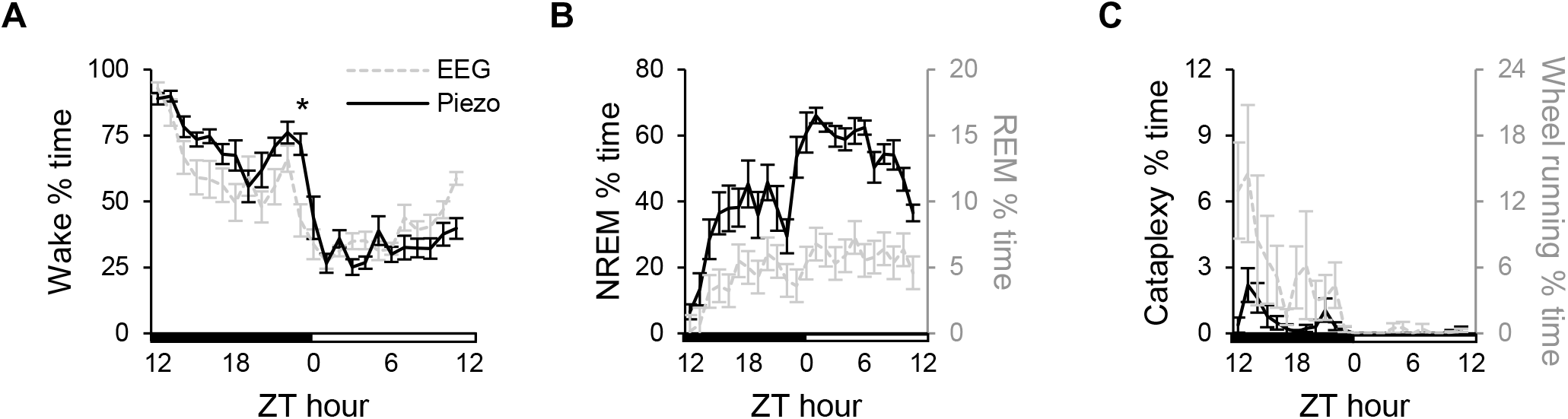
**Quantification of arousal states in narcoleptic mice by EEG/EMG recording.** Comparison of wake % per h in male orexin/ataxin-3 (Atax, *N* = 11) mice as measured by piezoelectric monitoring **(A, black line)** vs. manual scoring **(A, gray dashed line)** of EEG/EMG/video recordings from the same mice 8 weeks later. The % per h of NREM **(B, black line)**, REM **(B, gray dashed line)**, cataplexy **(C, black line)**, and wheel running **(C, gray dashed line)** as measured by manual scoring of EEG/EMG/video recordings. Two-way repeated-measures analysis of variance with Bonferroni’s multiple comparisons: * P < 0.05 vs. EEG/EMG/video.

Time spent in NREM and REM sleep (Figure 5B) and during cataplexy and wheel-running (Figure 5C) were quantified in these mice from the EEG/EMG with video recordings. During the dark period, 9 of the 11 mice (82%) exhibited ≥ 3 bouts of cataplexy, the typical criteria for inclusion in pharmacological studies on cataplexy (20). These mice exhibited 5.4 ± 0.9 bouts of cataplexy that each lasted 1.2 ± 0.2 min long for 4.7 ± 1.2 cumulative minutes during the dark period.

### Sleep/wake pharmacology using PiezoSleep

To validate piezoelectric monitoring of sleep/wake for the screening of wake-promoting compounds for narcolepsy, male Atax mice (*N* = 46) and WT mice (*N* = 60) were dosed with MOD (10, 30, 50, 100 mg/kg, 10 mL/kg) or vehicle orally at ZT5 when sleep propensity is high. As expected, MOD induced a dose-related inhibition of sleep for 3 h (30 and 50 mg/kg) and 6 h (100 mg/kg) post dosing vs. vehicle in WT (Figure 6A), *F*(24, 330) = 11.4, *P* < 0.0001. In Atax mice, MOD inhibited sleep for 2, 4, and at least 7 h post dosing at 30, 50, and 100 mg/kg, respectively, compared to vehicle (Figure 6B), *F*(24, 246) = 8.25, *P* < 0.0001. Although a dose-related response to MOD (10-100 mg/kg) was evident for the maximum sleep deficit in both Atax and WT mice (Figure 6C), *F*(4, 96) = 96.0, *P* < 0.0001 (main effect for dose), there was no interaction with genotype *F*(4, 96) = 1.63, *n.s*.

**Figure 6.**
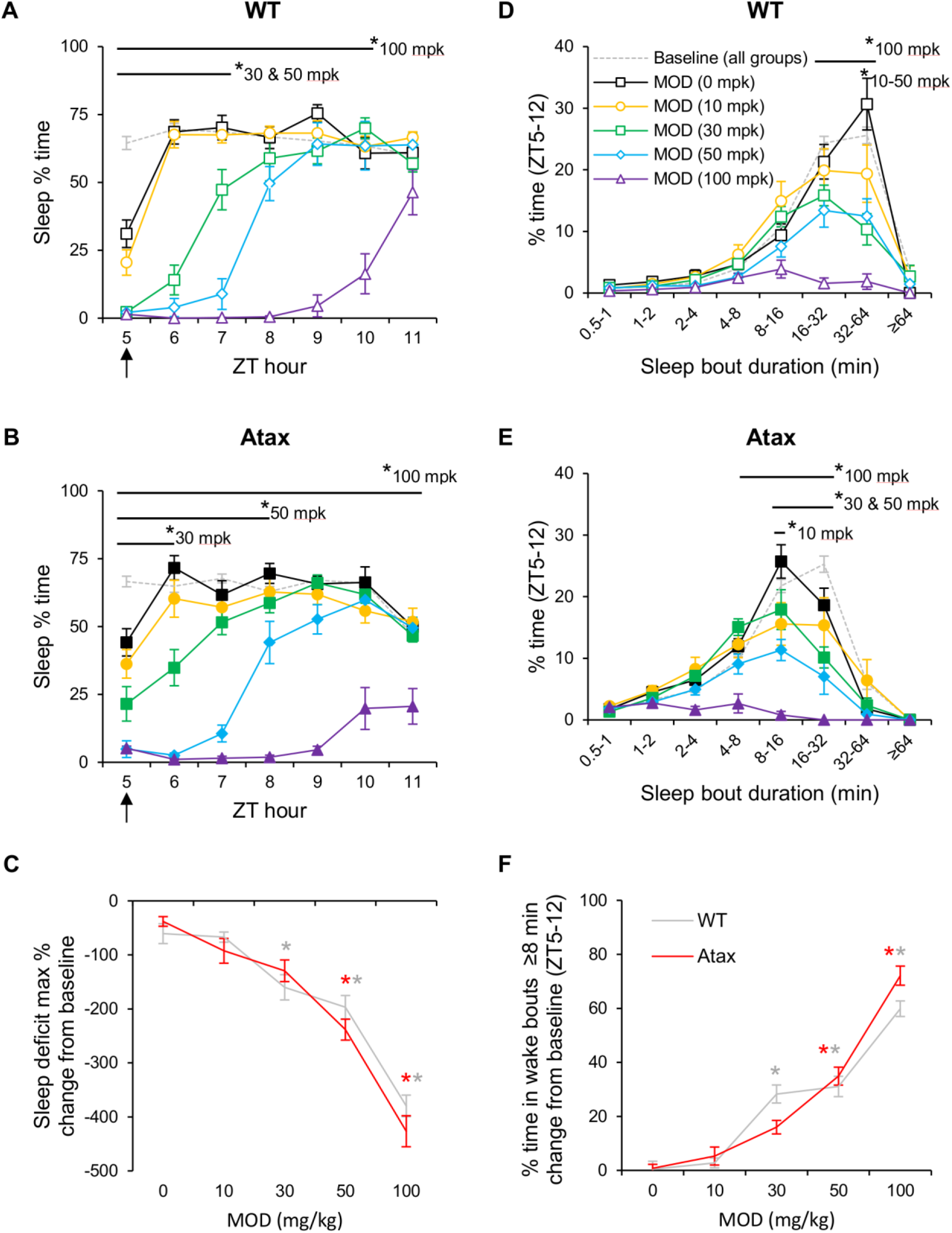
**Piezoelectric monitoring of sleep/wake in pharmacological studies.** Inhibition of sleep and promotion of wakefulness by modafinil (MOD) in orexin/ataxin-3 (Atax, **filled symbols, B, E** and **red lines C, F**, *N* = 9-10/group) vs. wild type (**open symbols, A, D** and **gray lines C, F**, *N* = 12/group) male mice as detected by PiezoSleep. Mice were administered MOD at 0 (vehicle, **black squares**), 10 (**orange circles**), 30 (**green squares**), 50 (**blue diamonds**), or 100 (**purple triangles**) mg/kg (mpk) at 5 h after lights on (zeitgeber time (ZT)5, **arrow**) after 30 h undisturbed baseline. Sleep % per h (**A-B**) and the percentage of time spent in sleep bout durations lasting from 0.5 to ≥ 64 min (**D-E**) for 7 h post dosing were plotted against values taken from undisturbed baseline (**gray dashed line**) for reference. The maximum sleep deficit © and the % of time spent in consolidated wake bouts ≥ 8 min for 7 h post dosing, expressed as the change from baseline (**F**), revealed dose-related wake-promotion by MOD (differences between genotypes, *n.s*.). Two-way analysis of variance with Bonferroni’s multiple comparisons: * P < 0.05 vs. vehicle for doses and at time or bout duration bin ranges as indicated by bars at top of graphs.

Modafinil fragmented sleep bouts in both mouse models. Under baseline and vehicle conditions during the last 7 h of the light period, WT mice spent most of their sleep time (26-31% of total time) in bouts that were 32-64 min long (Figure 6D). After MOD (10-100 mg/kg), the percentage of time spent in these long sleep bouts was reduced in dose-related manner, *F*(28, 385) = 5.38, *P* < 0.0001, and the distribution of time spent in the different SBD bins shifted such that more time was spent in shorter bouts than longer bouts (i.e., leftward shift of the peak), especially at the highest dose. Unlike WT mice, Atax mice in baseline and vehicle conditions spent 5-6% of the time in long sleep bouts ≥ 32 min, and most of their sleep time (22-25% of total time) was in bouts that were 8-32 min long (Figure 6E). Modafinil (10-100 mg/kg) further fragmented the sleep of Atax mice, as the percentage of time spent in sleep bouts 8-32 min long was reduced in a dose-related manner, *F*(24, 246) = 8.25, *P* < 0.0001. However, only the highest dose of MOD shifted the peak of the sleep bout distribution curve to the left toward shorter bouts. A dose-related increase in the percentage of time spent in wake bouts ≥ 8 min was evident after MOD (10-100 mg/kg) in WT and Atax mice (Figure 6F), *F*(4, 96) = 98.6, *P* < 0.0001 (main effect for dose), but the effect did not depend on genotype *F*(4, 96) = 2.87, *P* = 0.03, no significant contrasts per dose.

## Discussion

Piezoelectric monitoring of sleep/wake was used to identify key features of the narcoleptic phenotype that distinguish Atax mice by sleepiness severity, thus enabling rapid, non-invasive sorting of mice for further evaluation with EEG/EMG. The PiezoSleep narcolepsy screen was used to assign a WMS—the ratio of time spent in long WBD (≥ 32 min) to time in short SBD (< 8 min) during the first half of the dark period—to individual mice. The WMS with a cutoff value of 3.25 accurately differentiated genotype-confirmed Atax mice from WT controls. The WMS spanned a 15-fold difference between the most severely narcoleptic mice and their WT littermates. The PiezoSleep system was also demonstrated to detect changes in sleep/wake time and consolidation after administration of the wake-promoting therapeutic modafinil in normal and transgenic mice. Using this method, the efficacy of wake-promoting compounds can be determined in a 5-arm study with up to 60 mice in less than one week for high-throughput screening *in vivo*.

The PiezoSleep narcolepsy screen was designed to be a translational test of the ability to sustain long periods of wakefulness as is assessed in humans by the MWT and the MSLT. For the human tests, patients are provided 4-5 opportunities throughout the daytime to either nap within 20 min (MSLT) or to try to stay awake for 20-40 min (MWT) in the absence of alerting factors. The latency to polysomnographically defined sleep is measured for each test. A mean sleep onset latency ≤ 8 min with ≥ 2 sleep-onset REM periods (abnormal transition into REM sleep within 15 min of sleep onset) is a positive result on the MSLT and aids in the diagnosis of narcolepsy (21). A murine MSLT has been devised in which the latency to EEG/EMG-defined sleep is measured during four 20-min nap opportunities presented after 20 min of enforced wakefulness (27). The PiezoSleep narcolepsy screen provides a rapid, automated alternative to the murine MSLT and does not require the manipulation of sleep homeostatic processes to obtain the measurements. The 89% sensitivity and 92% specificity of the WMS on the PiezoSleep narcolepsy screen for mice is reasonably comparable to that of sleep latency on the MSLT (95% sensitivity and 73% specificity) and MWT (84% sensitivity and 98% specificity) for humans (31). The translational utility of the PiezoSleep phenotyping assay is limited by the inability of the current version of the algorithm to distinguish between NREM and REM sleep. Perhaps a new algorithm under development (32) will permit quantification of short REM sleep latency into the assay as a murine equivalent of sleep-onset REM periods.

The PiezoSleep narcolepsy screen can be used to prescreen narcoleptic mice to identify those with the most fragmented sleep/wake for inclusion in resource-intensive EEG/EMG studies. The WMS clearly differentiated narcoleptic mice with highly fragmented sleep/wake from WT littermates that were able to spontaneously sustain long bouts of wakefulness from ZT12-18 (Figure 2). It is unknown if the wake bouts in Atax mice terminated because of sleep onset or from the intrusion of cataplexy, which may register on the PiezoSleep assay as sleep due to the similar motor physiology between REM sleep and cataplexy. Nevertheless, this distinction does not detract from the utility of the PiezoSleep narcolepsy screen to differentiate Atax mice by severity of sleep/wake fragmentation. Out of the 37 Atax mice screened for inclusion in the EEG/EMG study, 16 mice with severe symptoms were identified through rank ordering of the WMS. These mice scored 4.7x lower on the PiezoSleep narcolepsy screen than the remainder of the Atax mice, and only 2 out of the final 11 included in analysis (18%) had < 3 cataplexy bouts. Compared to an unpublished survey of 39 Atax mice in which 34% of the mice fell below this study inclusion criteria (20), the current study represents a 16% improvement in yield. However, it is difficult to determine if the yield differences are due to the new sorting methodology or to differences between colonies and recording environments. The reason for high interindividual differences in cataplexy expression between Atax mice remains unclear, but appears to reflect the variation observed in human narcolepsy (20). Determination of whether the WMS on the PiezoSleep narcolepsy screen correlate with the severity of cataplexy would require EEG/EMG measurement from mice across a broader range of symptom severity scores and was beyond the scope of the current study. Comparison of EEG/EMG-defined arousal states between mice with low and high WMS identified by piezoelectric monitoring would be an important next step in the validation of the PiezoSleep narcolepsy screen.

Some differences between studies in the comparison of piezoelectric monitoring of sleep/wake and polysomnography are worth noting. Previous studies have validated the PiezoSleep sleep/wake classifier against manually scored EEG/EMG signals that had been simultaneously recorded from the same mice (across 4 strains) for direct epoch-to-epoch comparison (9, 15). The current analysis in Atax mice does not permit this level of scrutiny, as the PiezoSleep and EEG recordings were staged in 2 and 10 second epochs, respectively, and occurred 8 weeks apart in different environments (e.g., the EEG recordings occurred in standard cages equipped with running wheels in a sound-attenuated recording chamber). Wakefulness over 24 h was 3.6% higher after piezoelectric monitoring than after EEG/EMG recording in the current study, vs. 2.3% lower as previously reported (15). In addition to the caveats between methods as noted above, the discrepancy could be due to the different mouse strains studied (Atax vs. CFW Swiss Webster). It is unlikely that the breathing patterns of Atax mice would cause the PiezoSleep assay to misconstrue sleep as quiet wakefulness, and thus overinflate the measurement of time awake compared to EEG/EMG in the current study. Piezoelectric monitoring did not detect more wakefulness in Atax mice than in WT (Figure 2A). Although orexin knockout mice show an attenuated respiratory response to hypercapnia during wakefulness compared to WT, breath rate and tidal volume in normal room air are indistinguishable between genotypes across arousal states (33, 34). The differences in sleep/wake as determined by EEG/EMG and piezoelectric monitoring in the current study are likely due to the different environments used for recording. Future studies are needed in which simultaneous piezoelectric EEG/EMG recordings are collected during the dark period from narcoleptic mice to assess cataplexy detection using an unsupervised algorithm that has been tuned to extract features of REM sleep (32) to determine if cataplexy can be identified by piezoelectric monitoring.

Perhaps more noteworthy than the small discrepancies between polysomnography and PiezoSleep monitoring in the current study was the consistency in the pattern of sleep/wake between the two recording methods (Figure 5A). The hour-by-hour increase and decrease in wake time after EEG/EMG recording paralleled the pattern observed after piezoelectric monitoring, despite the 8-week difference in recording dates. The hour immediately prior to lights-on was the only time point in which PiezoSleep registered more wakefulness in Atax mice than polysomnography (i.e., At ZT23, Atax mice slept more as determined by polysomnography than they did 8 weeks prior as measured with PiezoSleep). This time point was the same ZT in which Atax mice slept more than WT mice after piezoelectric monitoring of both genotypes. The increased sleep observed in Atax mice at the end of the dark period may reflect progressive worsening of EDS over time in this model. The differences in sleep/wake measurement between genotypes and between methods at ZT23 are less likely due to reduced accuracy of PiezoSleep at that time point, as could be speculated, because the variance was similar at that hour between the different runs and between other ZT hours. It is also not clear what physiological or behavioral factors would differentially impact the detection of respiration rates and gross motor activity at ZT23 vs. other times of day.

In the current study, piezoelectric measurement of sleep/wake was evaluated as a method to screen wake-promoting compounds rapidly and at scale for the development of therapeutics for narcolepsy. Dosing WT mice at ZT5 was chosen as the first pass of screening using MOD because of the similarities between Atax and WT mice in the high amount of sleep per hour (Figure 2A) and in the very short WBD (Figure 2C) during the light period. Modafinil (50 mg/kg) with total brain exposure of 33.7 μM approximated the ED_50_ in both WT and Atax mice in terms of the time course of sleep inhibition, the maximum sleep deficit induced, and the consolidation of wakefulness (Figure 6) and is recommended as the positive control for screening the wake-promoting efficacy of compounds. Using the PiezoSleep assay, a total of 106 mice in two cohorts were screened across 4 concentrations of MOD vs. vehicle in two weeks, representing substantial improvement in efficiency over conventional sleep/wake bioassays (22, 30). Future studies are needed to validate the PiezoSleep assay for pharmacological studies at different times of day and with hypnotics. Of particular interest would be to test the hypothesis that wake-promoting therapeutics administered at ZT12 would rescue the narcoleptic phenotype of Atax mice as measured by the WMS on the PiezoSleep narcolepsy screen.

Piezoelectric monitoring of sleep/wake offers advantages over video-based methods, even those that claim to discriminate NREM and REM sleep (35) for several reasons. Because PiezoSleep does not depend on a visual signal, it permits mice to sleep while burrowed in nesting material for assessment of more ecologically relevant behavior. The highly sensitive piezoelectric sensor captures a detailed respiratory measure that enables it to distinguish quiet, resting wakefulness from sleep using relevant physiology (15). To the best of our knowledge, sleep state discrimination based on postural changes in the aspect ratio of the body on video recordings (35) has not been replicated nor validated in mouse models that have been reported to feature obesity, such as Atax mice (18, 36).

## Conclusions

The non-invasive, unsupervised piezoelectric monitoring of sleep/wake is a new tool for efficacy studies with arousal state endpoints and can be used to solve two problems in the preclinical development of wake-promoting therapeutics for narcolepsy. First, the PiezoSleep narcolepsy screen can be used to determine a WMS that quantifies sleepiness severity and can be used to improve phenotypic sorting for resource intensive EEG/EMG studies. Second, PiezoSleep can be used to rapidly quantify sleep/wake time and consolidation for high-throughput screening *in vivo* for drug discovery.

## List of Abbreviations

ANOVA: analysis of variance
Atax: *orexin/ataxin*-3
AUC: area under the curve
Cmax: maximum concentration
EDS: excessive daytime sleepiness
EEG: electroencephalography
EMG: electromyography
f: fornix
ic: internal capsule
LD12:12: 12 h light period, 12 h dark period
MOD: modafinil
MSLT: multiple sleep latency test
MWT: maintenance of wakefulness test
NREM: non-rapid-eye-movement
opt: optic tract
PBS: phosphate-buffered saline
PO: *per os*
REM: rapid-eye-movement
RM ANOVA: repeated-measures analysis of variance
ROC: receiver operator characteristic
SBD: sleep bout duration
SC: subcutaneous
S.E.M.: standard error of the mean
mpk: mg per kg
MRT: mean residence time
T_1/2_: half life
WBD: wake bout duration
WT: wild type
ZT: zeitgeber time
3V: third ventricle

## Declarations

### Ethics approval and consent to participate and permissions

This research did not involve human subjects, human material, nor human data; consent to participate is not applicable. All experimental procedures on animals were approved by the Institutional Animal Care and Use Committee (IACUC Protocol No. IP-008 and IP-010) at Reset Therapeutics and were conducted in accordance with the principles set forth in the *Guide for Care and Use of Laboratory Animals of the National Institute of Health*. All experiments were in compliance with the ethical guidelines from the International Council for Laboratory Animal Science and in accordance with the Basel Declaration.

### Consent for publication

Not applicable

### Availability of data and materials

The datasets generated and/or analyzed during the current study are not available in public repositories due to ownership by private enterprise, but they can be made available to interested parties on reasonable request.

### Authors’ contributions

S.W.B. designed and managed the efficacy studies, analyzed the data, prepared the figures and wrote the manuscript. J.D.S. designed and managed the PK study, prepared the table, formulated the compound, scored the data and reviewed the manuscript. A.L., N.K., and P.S. executed the studies, performed surgeries, scored the data and reviewed the manuscript. K.D. and B.O.H. provided the PiezoSleep system, created custom analysis tools, and reviewed the manuscript. R.B. and P.S.H. conceptualized the research goals, supervised the team, acquired funding and resources, and reviewed the manuscript.

## Acknowledgements

We thank Drs. Thomas Scammell (Harvard University) and Takeshi Sakurai (University of Tsukuba) for the donation of the orexin/ataxin-3 mice to Jackson Labs, Dr. Orsolya Kiraly (The Jackson Laboratory) for performing the histology, and Tod Steinfeld and Dr. Bruce Clapham (Reset Therapeutics) for helpful discussions.

## Authors’ information (optional)

Not applicable

